# Global and regional white matter development in early childhood

**DOI:** 10.1101/524785

**Authors:** Jess E. Reynolds, Melody N. Grohs, Deborah Dewey, Catherine Lebel

**Author notes:** Corresponding Author: Catherine Lebel, Room B4-513, Alberta Children’s Hospital 28 Oki Drive NW, Calgary, Alberta, Canada, T3B6A8.

## Abstract

White matter development continues throughout childhood and into early adulthood, but few studies have examined early childhood, and the specific trajectories and regional variation in this age range remain unclear. The aim of this study was to characterize developmental trajectories and sex differences of white matter in typically developing young children. Three hundred and ninety-six diffusion tensor imaging datasets from 120 children (57 male) aged 2-8 years were analyzed using tractography. Fractional anisotropy (FA) increased and mean diffusivity (MD) decreased in all white matter tracts by 5-15% over the 6-year period, likely reflecting increases in myelination and axonal packing. Males showed steeper slopes in a number of brain areas. Overall, early childhood is associated with substantial development of all white matter and appears to be an important period for the development of occipital and limbic connections, which showed the largest changes. This study provides a detailed characterization of age-related white matter changes in early childhood, offering baseline data that can be used to understand cognitive and behavioural development, as well as to identify deviations from normal development in children with various diseases, disorders, or brain injuries.

## 1. Introduction

Rapid brain microstructural maturation occurs during the first years of life, with remodelling processes continuing at a slower pace into adulthood (Dubois et al., 2014; Lebel et al., 2008b; Simmonds et al., 2014; Taki et al., 2013; Tamnes et al., 2009). Diffusion MRI enables in vivo exploration of white matter microstructural development in infants, young children, adolescents, and adults. Limited research, however, has detailed brain white matter development over the preschool period (Brown and Jernigan, 2012), in part due to the difficulties associated with MRI scanning of young children (e.g. motion). The dynamic changes in behavior, cognitive abilities, and emotional regulation that are characteristic of early childhood (2-8 years) make this an age range of particular interest.

Diffusion MRI studies show rapid white matter development in the first two years of life (increases in fractional anisotropy [FA], reductions in mean diffusivity [MD]) that follows posterior to anterior and central to peripheral patterns of development, and reflects ongoing myelination (Dubois et al., 2006, 2008; Krogsrud et al., 2016; Qiu et al., 2015). From late childhood (~6 years) to young adulthood, slower increases of FA and decreases of MD are most likely associated with increases of axonal packing/density (Hermoye et al., 2006; Lebel et al., 2008b; Taki et al., 2013). Fronto-temporal fiber connections (e.g. uncinate, cingulum) demonstrate the most protracted development, continuing into the late 20s and 30s (e.g. Lebel et al., 2008b; Simmonds et al., 2014).

Differences between males and females in the prevalence and timing of onset of neurodevelopmental disorders (Giedd et al., 2012) underscore the importance of considering sex differences in brain development. There is strong evidence for sexual dimorphism in brain volume growth. Studies consistently demonstrate that males have larger brains across the lifespan, and that females reach peak brain volumes approximately 1.5 years earlier than males (Lenroot et al., 2007). Findings of sex differences in diffusion studies have been much more mixed. A small number of studies in later childhood and adolescence suggest earlier white matter development in girls (Asato et al., 2010; Bava et al., 2011; Clayden et al., 2012; Seunarine et al., 2016; Simmonds et al., 2014; Wang et al., 2012), but it is not clear when trajectories diverge.

Our current understanding of typical brain development in early childhood is based largely on small cross-sectional studies (Huang et al., 2006), or studies of larger age-ranges that include only a small number of young children (Genc et al., 2017; Hagmann et al., 2010; Krogsrud et al., 2016; Löbel et al., 2009; Moon et al., 2011; Sadeghi et al., 2015; Schneider et al., 2004), which limits a detailed understanding of development during this period. While there is increasing interest in early neuroimaging biomarkers of developmental disorders (e.g., autism (Hazlett et al., 2017; Varcin and Jeste, 2017)), the lack of a detailed understanding of early typical brain development hinders our ability to identify atypical trajectories. A more detailed understanding of development trajectories, and knowledge of subtle variations of brain maturation during this period, including the specific timing of changes, would provide baseline data that could be used to examine deviations from typical development, and relationships with cognitive abilities and behavior. The primary aim of this study was to examine white matter development trajectories within and across young children (2–8 years), including regional variation and sex differences. We used diffusion tensor imaging (DTI) to measure FA and MD globally and in ten major white matter tracts on 396 scans from 120 typically developing children. We hypothesized that FA would increase and MD would decrease at decelerating rates across early childhood. Based on infant literature (Dubois et al., 2014; Paydar et al., 2014), we hypothesized initially higher FA and lower MD within motor (e.g. corpus callosum, pyramidal) tracts. Furthermore, we hypothesized slower and later maturation of fronto-temporal fiber bundles, which have been demonstrated to have a more protracted development course in adolescents and adults (Lebel et al., 2012).

## 2. Material and Methods

### 2.1 Participants

This study included 396 datasets on 120 typically developing children (57 male). Participants were aged 1.95-6.97 years at intake (mean = 4.04 ± 1.07 years) and were recruited from the Calgary area and from the ongoing prospective Alberta Pregnancy Outcomes and Nutrition (APrON) Study (Kaplan et al., 2014). Children were asked to return approximately semi-annually, though not all returned for each visit. Average time between visits was 7.99 ± 4.63 months. Data analyzed here includes 41 children with only one scan time point, 20 children with two scans, 12 with three scans, 11 with four scans, 13 with five scans, 9 with six scans, 12 with seven scans, 1 with twelve scans, and 1 with twenty scans (Figure 1). All children were free from diagnosed neurological, genetic, and neurodevelopmental (e.g. ADHD, ASD) disorders and were born ≥ 36 weeks gestation. Both maternal and paternal post-secondary education ranged from 2 to 13 years (all parents completed high school), with means of 5.78 ± 2.76 years (n = 101 parents completed the questionnaire) and 4.88 ± 2.72 years (n = 101), respectively. The majority of the sample was Caucasian (93% of mothers and 89% of fathers). Other self-reported ethnicities included Asian/Pacific Islander, African, Filipino, Latino/Hispanic, and Multiracial. Annual household income ranged from <$25,000 to >$175,000; the median income range was the $125,000 - $149,999 bracket. Child sex and parental education, ethnicity, and household income at intake did not differ between participants with 1-2 scans compared to those with ≥4 scans (*p* > 0.05). Parental/guardian consent and child assent were obtained for each subject. The University of Calgary Conjoint Health Research Ethics Board (CHREB) approved this study (REB13-0020).

**Figure 1:**
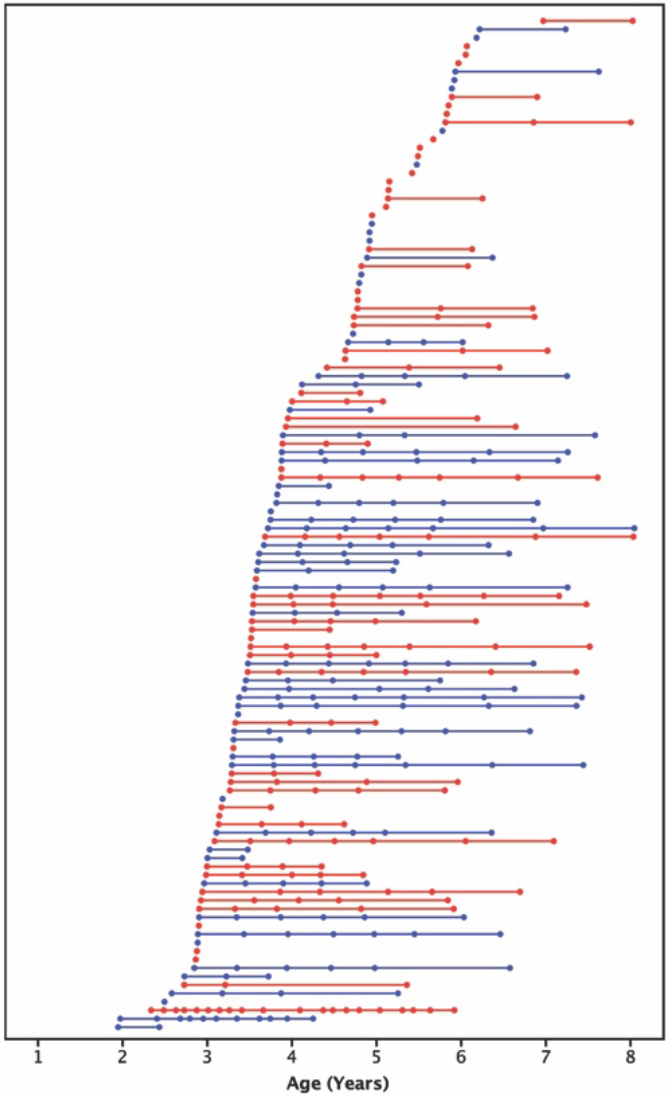
Age at scans for all subjects (females are shown in red, males shown in blue). Each of the 396 scans is represented by a circle; each of the 120 subjects is shown in a different row with their scans connected by a straight line.

### 2.2 MRI Image acquisition

All imaging was conducted using the same General Electric 3T MR750w system and 32-channel head coil (GE, Waukesha, WI) at the Alberta Children’s Hospital, Calgary, Canada. Children were scanned either while awake and watching a movie, or while sleeping without sedation. Prior to scanning, parents were provided with detailed information on MRI procedures and were given the option to complete a practice MRI session in an MRI simulator to familiarize their child with the scanning environment, and/or to make use of a take home pack with information on MRI scanning (e.g., noise recordings (Thieba et al., 2018)).

Whole-brain diffusion weighted images were acquired using single shot spin echo echo-planar imaging sequence with: 1.6 x 1.6 x 2.2 mm resolution (resampled on scanner to 0.78 x 0.78 x mm), full brain coverage (matrix size: 256 x 256 x 54), FOV = 200mm x 200mm, TR = 6750 ms; TE = 79 ms, flip angle = 90°, anterior-posterior phase encoding, 30 gradient encoding directions at b=750s/mm^2^, and five interleaved images without gradient encoding at b=0s/mm^2^, for a total acquisition time of 4:03 minutes.

### 2.3 DTI data processing and semi-automated fiber tracking

DTI data was visually quality checked and volumes with artifacts or motion corruption were removed (e.g. venetian blinds, vibration; See Supplementary Figures 1-6 for examples of artifacts, and representative diffusion weighted and b0 images, and image maps). Entire datasets were excluded if fewer than 18 high quality diffusion weighted and 2 b0 volumes remained. Of participants included in the analyses reported here, the average number of volumes remaining was 27.5 ± 2.9 (range = 18-30) diffusion weighted and 4.8 ± 0.5 (range = 2-5) b0 images (Figure 2). There was a weak positive correlation between age and number of volumes remaining (*r* = 0.251, *p* <0.001). Volumes removed were not related to child sex, or parental education (*p* > 0.05). Data was then pipelined through ExploreDTI V4.8.6 (Leemans et al., 2009) for preprocessing. Corrections for signal drift, Gibbs ringing (non-DWIs), subject motion, and eddy current distortions were performed (Leemans and Jones, 2009).

**Figure 2.**
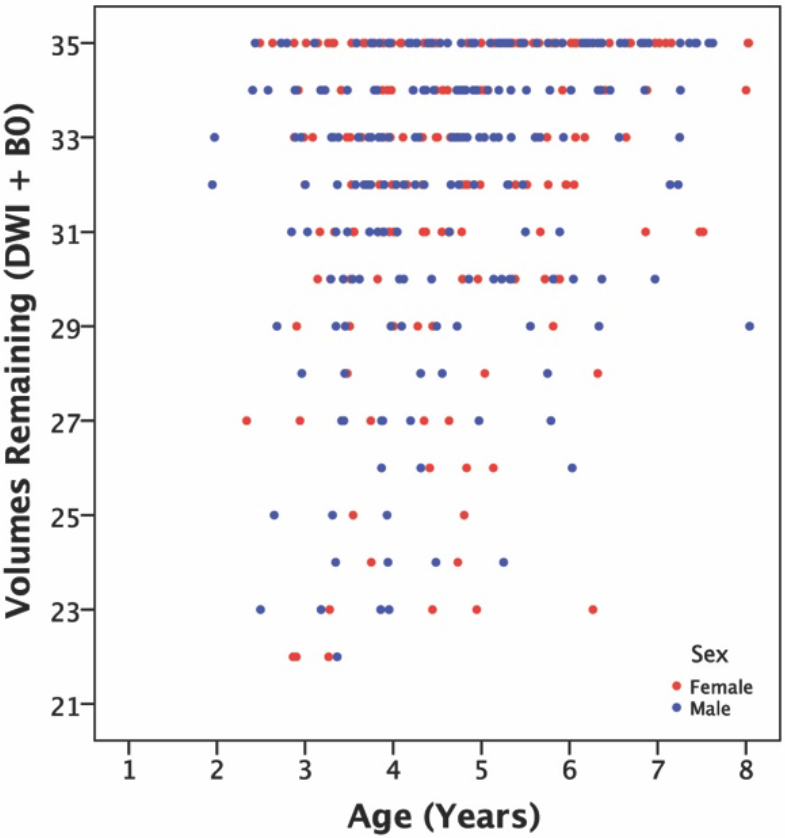
Number of volumes remaining for each scan following bad volume removal, shown by age and sex (males = blue, females = red).

Semi-automated deterministic streamline tractography was used to delineate 10 major white matter tracts: the pyramidal tract, corpus callosum (genu, body, splenium), cingulum, fornix, inferior fronto-occipital fasciculus (IFO), inferior longitudinal fasciculus (ILF), superior longitudinal fasciculus including arcuate (SLF), and uncinate fasciculus (UF) (region of interest semi-automated tractography guides can be found at: https://doi.org/10.6084/m9.figshare.7603271.v1). The minimum FA threshold was set to 0.20 to initiate and continue tracking, and the angle threshold set to 30º. The most representative scan (a 3.68 year old female) was selected in FSL (Andersson et al., 2007; Jenkinson et al., 2012; Smith et al., 2004; Woolrich et al., 2009) for use as a target in semi-automated tractography. To identify the target scan, all scans collected prior to September 2017 (when semi-automated tractography commenced) with ≥34 high quality volumes were input, and each FA map was nonlinearly registered to every other FA map. The most representative scan (target scan) was selected as the FA map that required the least amount of warping when being registered to all the other FA maps. Using a priori information on tract location (Lebel et al., 2008b; Wakana et al., 2004), inclusion and exclusion regions of interest (ROI) were drawn on the target participant. The inclusion and exclusion were formulated and tested together by two researchers (JR, MG) using an iterative testing and verification procedure. All scans were normalized to the target scan using a non-affine transformation, and the inverse normalization (i.e., target to each scan) was calculated. Inclusion and exclusion regions from the target were the applied to each scan using the inverse normalization (Lebel et al., 2008a). All tracts were manually quality checked and additional exclusion ROIs were drawn as required on a small number of tracts to remove spurious fibers. The operators were blinded to participant ID.

### 2.4 White matter fiber tract and global white matter diffusion measurements

Mean FA and MD measures were extracted from all tracts using ExploreDTI. Eigenvalues of the tracts were also extracted to calculate axial diffusivity (AD) and radial diffusivity (RD). Where appropriate (i.e., for all tracts except the corpus callosum and fornix), values were calculated separately for left and right hemisphere tracts and subsequently averaged. FA and MD models for left and right hemisphere tracts are presented in supplementary material (Supplementary Tables 3-4). In addition, mean MD and FA across all white matter fibers were calculated to provide global white matter measures.

### 2.5 Statistics and curve fitting

Statistics were performed using RStudio version 1.1.453 (RStudio Team, 2016), and the ‘lme4’ (Bates et al., 2015), and ‘lmerTest’ (Kuznetsova et al., 2017) packages. Linear mixed models were run to determine the developmental trajectories (rates of change) of FA and MD for each tract and global white matter metrics. MD values were scaled by 1000 to bring them to a similar scale as other measures for analysis; reported values were scaled back to reflect actual values for ease of interpretation. Linear mixed models with linear (y = age + age*sex + (1|subject)) and quadratic (y = age + age^2^ + age*sex + (1|Subject)) terms were modelled, with age, sex, and age*sex interactions modelled as fixed predictors and subject modelled as a random factor. Restricted maximum likelihood (REML) was set to false, and the Bayesian information criterion (BIC) for each model was compared for model selection, whereby smaller and negative values were considered the best fit. To compare tract development, both absolute and percent change (absolute change divided by 2-year predicted value) from 2.0-8.0 years were calculated (based on predicted group values at 2 and 8 years). Rate of development for each tract was compared based on slope and absolute change across the early childhood period. Significance level was set at *p* < 0.05; multiple comparisons were corrected within each factor (e.g., age) of the models (2 diffusion metrics: FA and MD, 10 tracts: 0.05/20 = 0.0025). Both corrected (*p* < 0.0025) and uncorrected (*p* < 0.05) level results are reported. Linear mixed models were also run including the number of remaining volumes, white matter volume, and the tract volume as fixed effects.

### 2.6 Data and code availability statement

A guide for regions of interest used in the semi-automated tractography can be found at: https://doi.org/10.6084/m9.figshare.7603271.v1. Data and code used in this study will be made available upon request to the corresponding and/or first author.

## 3. Results

### 3.1 Trajectories of white matter development during the early childhood period

All tracts, as well as global means (Figure 3), showed significant age-related increases of FA and decreases of MD, except fornix MD (Tables 1-2, Figures 4-5). For FA, the genu, splenium, fornix, pyramidal tract, and uncinate fasciculus demonstrated linear changes; the body of the corpus callosum, cingulum, IFO, ILF, SLF, and global FA demonstrated quadratic development. For MD, the splenium, and uncinate showed linear decreases, while the body, genu, cingulum, IFO, ILF, pyramidal, SLF, and global MD demonstrated quadratic declines.

**Table 1.** Linear regression parameters for FA. Global FA is presented first, followed by tracts ordered by percent change from 2-8 years (lowest to highest). Sex term represents boys: Positive sex coefficients indicate higher FA in boys; Positive age*sex interaction coefficients indicate more positive slopes in boys (i.e., faster increases of FA). Significance levels presented as *** <0.001, ** <0.0025 (Bonferroni corrected), * <0.05.

| Tract | Intercept (0 years) ± SE | Linear age term ± SE ($10^{-2}$) | Quadratic age term ± SE ($10^{-2}$) | Sex ± SE ($10^{-2}$) | Age*Sex ± SE ($10^{-2}$) | 2 years (predicted) | Absolute change 2-8 years | % change 2-8 years |
| --- | --- | --- | --- | --- | --- | --- | --- | --- |
| Global FA | 0.41±0.004*** | 1.09±0.136*** | -0.05±0.013** | 0.22±0.266 | -0.06±0.043 | 0.43 | 0.035 | 8.01 |
| Splenium (CC) | 0.51±0.003*** | 0.55±0.062*** | NA | 0.02±0.484 | -0.03±0.083 | 0.53 | 0.032 | 6.11 |
| Pyramidal | 0.44±0.003*** | 0.77±0.050*** | NA | 1.13±0.398* | -0.15±0.069* | 0.46 | 0.042 | 9.02 |
| Body (CC) | 0.43±0.008*** | 1.85±0.304*** | -0.12±0.029*** | -0.42±0.542 | 0.07±0.096 | 0.46 | 0.043 | 9.31 |
| Genu (CC) | 0.44±0.004*** | 0.78±0.075*** | NA | 1.18±0.577* | -0.09±0.104 | 0.46 | 0.044 | 9.56 |
| SLF | 0.38±0.006*** | 1.38±0.205*** | -0.07±0.020*** | 0.04±0.416 | -0.11±0.065 | 0.40 | 0.040 | 9.95 |
| Uncinate | 0.35±0.003*** | 0.70±0.056*** | NA | -0.21±0.456 | 0.02±0.077 | 0.37 | 0.043 | 11.70 |
| Fornix | 0.34±0.005*** | 0.60±0.102*** | NA | -0.93±0.739 | 0.22±0.142 | 0.35 | 0.043 | 12.07 |
| Cingulum | 0.35±0.008*** | 1.75±0.307*** | -0.09±0.030** | 0.22±0.576 | 0.003±0.097 | 0.39 | 0.050 | 13.02 |
| IFO | 0.37±0.006*** | 1.94±0.220*** | -0.10±0.021*** | -0.05±0.423 | -0.01±0.069 | 0.40 | 0.058 | 14.24 |
| ILF | 0.35±0.007*** | 2.51±0.253*** | -0.15±0.024*** | -0.28±0.477 | -0.06±0.080 | 0.39 | 0.058 | 14.93 |

**Table 2.** Linear regression parameters for MD (mm^2^/s). Global MD is presented first, followed by tracts ordered by percent change from 2-8 years (lowest to highest). Sex term represents boys: Positive sex coefficients indicate higher MD in boys; Negative age*sex interaction coefficients indicate more negative slopes in boys (i.e., faster decreases of MD). Significance levels presented as *** <0.001, ** <0.0025 (Bonferroni corrected), * <0.05.

| Tract | Intercept (0 years) ± SE ($10^{-4}$) | Linear age term ± SE ($10^{-4}$) | Quadratic age term ± SE ($10^{-4}$) | Sex ± SE ($10^{-4}$) | Age*Sex ± SE ($10^{-4}$) | 2 years (predicted) ($10^{-4}$) | Absolute change 2-8 years ($10^{-4}$) | % change 2-8 years |
| --- | --- | --- | --- | --- | --- | --- | --- | --- |
| Global MD | 9.95±0.071*** | -0.356±0.027*** | 0.023±0.003*** | 0.163±0.049** | -0.011±0.008 | 9.40 | -0.80 | -8.51 |
| Splenium (CC) | 9.15±0.079*** | -0.071±0.014*** | NA | 0.214±0.109 | -0.014±0.020 | 9.10 | -0.47 | -5.14 |
| Pyramidal | 9.23±0.079*** | -0.292±0.030*** | 0.019±0.003*** | 0.056±0.052 | 0.007±0.009 | 8.76 | -0.58 | -6.65 |
| Cingulum | 9.73±0.125*** | -0.399±0.048*** | 0.030±0.005*** | 0.157±0.082 | -0.013±0.015 | 9.12 | -0.65 | -7.18 |
| Fornix | 14.49±0.307*** | -0.098±0.054 | NA | 1.190±0.424* | -0.169±0.074* | 14.72 | -1.10 | -7.44 |
| SLF | 9.32±0.086*** | -0.340±0.032*** | 0.024±0.003*** | 0.260±0.061*** | -0.024±0.010* | 8.84 | -0.69 | -7.84 |
| Uncinate | 9.81±0.052*** | -0.138±0.009*** | NA | 0.036±0.071 | -0.005±0.013 | 9.55 | -0.84 | -8.80 |
| Body (CC) | 10.48±0.142*** | -0.447±0.054*** | 0.031±0.005*** | 0.280±0.098* | -0.025±0.017 | 9.82 | -0.90 | -9.15 |
| IFO | 10.05±0.102*** | -0.313±0.039*** | 0.017±0.004*** | 0.125±0.070 | -0.005±0.012 | 9.55 | -0.89 | -9.29 |
| ILF | 10.29±0.118*** | -0.488±0.045*** | 0.033±0.004*** | 0.371±0.079*** | -0.033±0.014* | 9.60 | -1.04 | -10.79 |
| Genu (CC) | 11.16±0.212*** | -0.456±0.082*** | 0.024±0.008* | -0.122±0.132 | 0.008±0.0255 | 10.29 | -1.28 | -12.39 |

**Figure 3.**
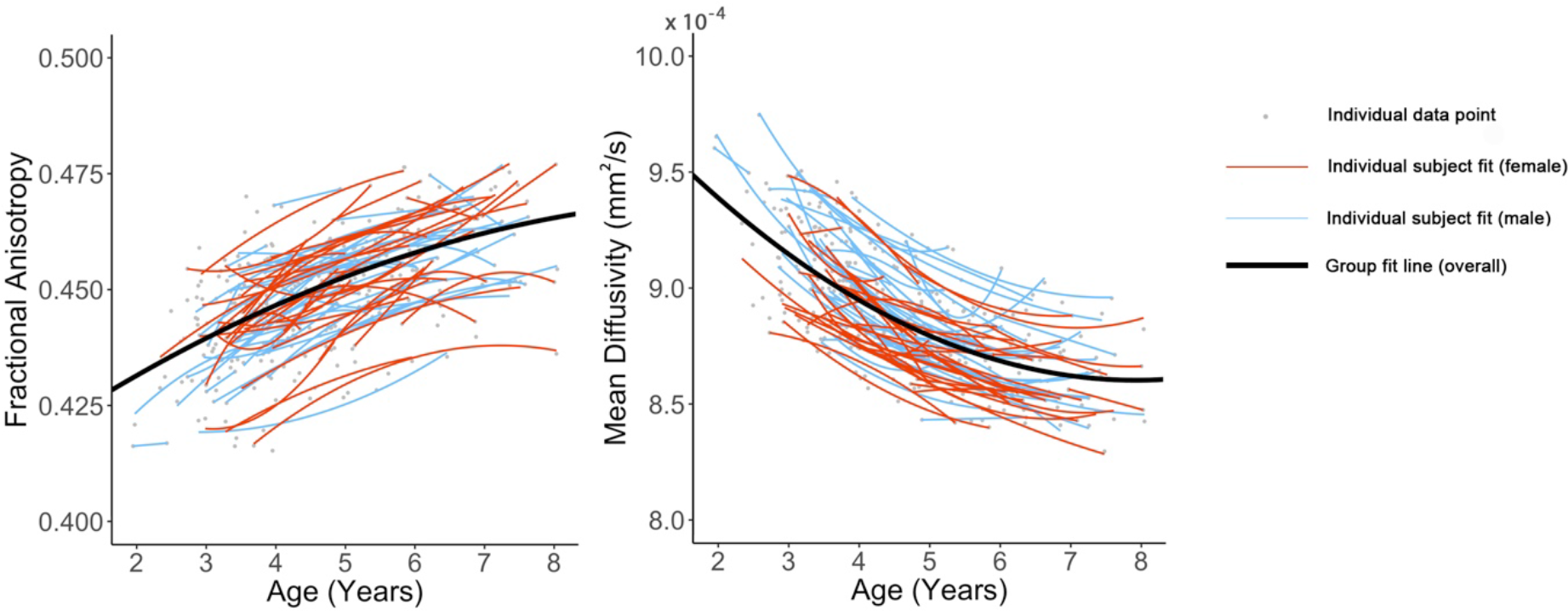
Trends of global mean FA and MD. Best fit lines for the entire dataset are shown in black (where there was no sex effect) or in blue and red separately for males and females (where the sex*age interaction was significant). Best fit lines for each individual subject are shown with thinner red (girls) and blue lines (boys). Individual data points are shown in grey.

**Figure 4.**
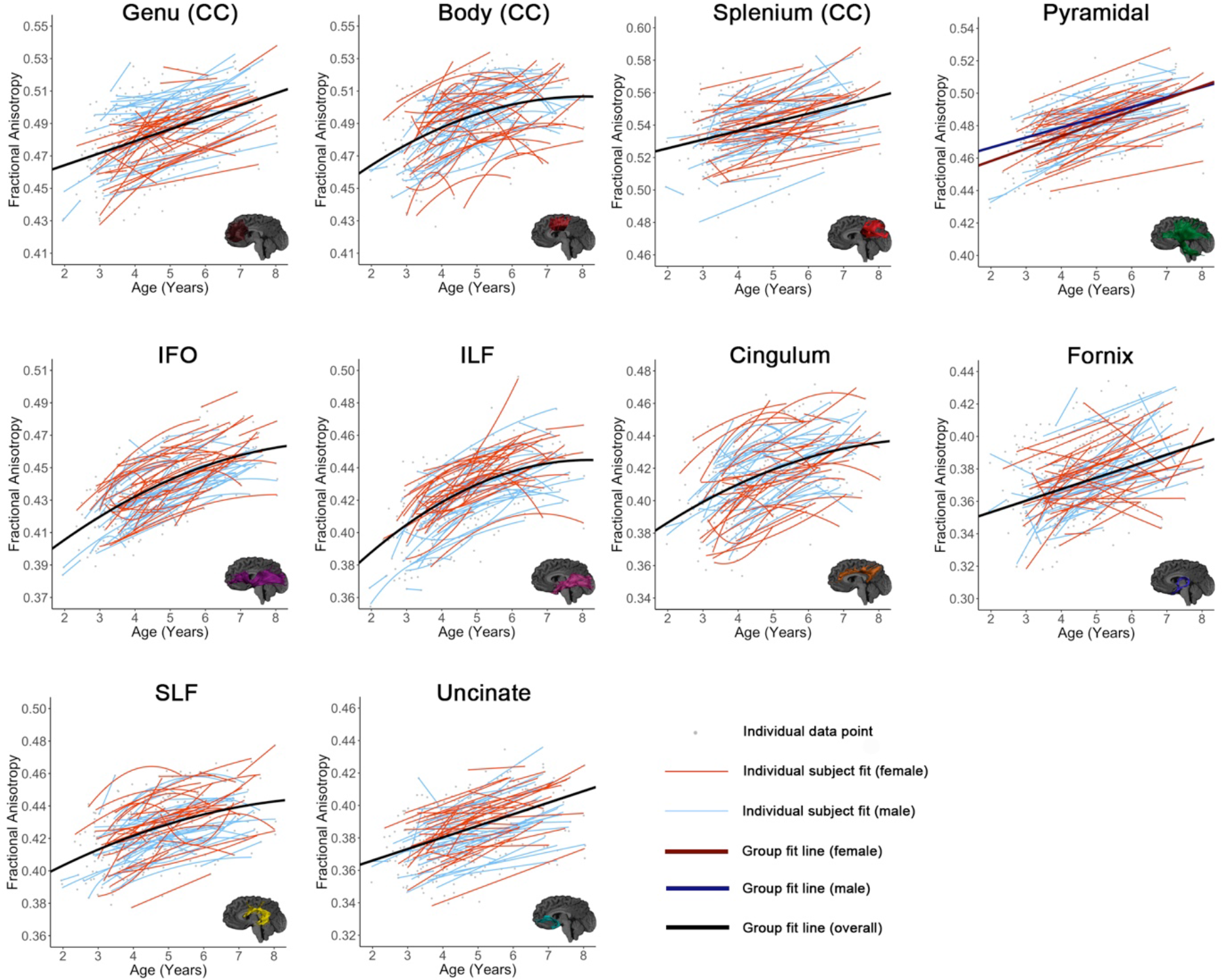
Relationships between age and FA are shown for all tracts. Tracts are grouped by high initial FA and slow development (row 1), low initial FA and rapid maturation (row 2), and low initial FA and slower development (row 3). Best fit lines for the entire dataset are shown in black (where there was no significant age*sex interaction) or red and blue (separated for girls and boys, where there was a significant interaction). Best fit lines for each individual subject are shown in thinner red (girls) and blue (boys) lines. Individual data points are shown in grey.

**Figure 5.**
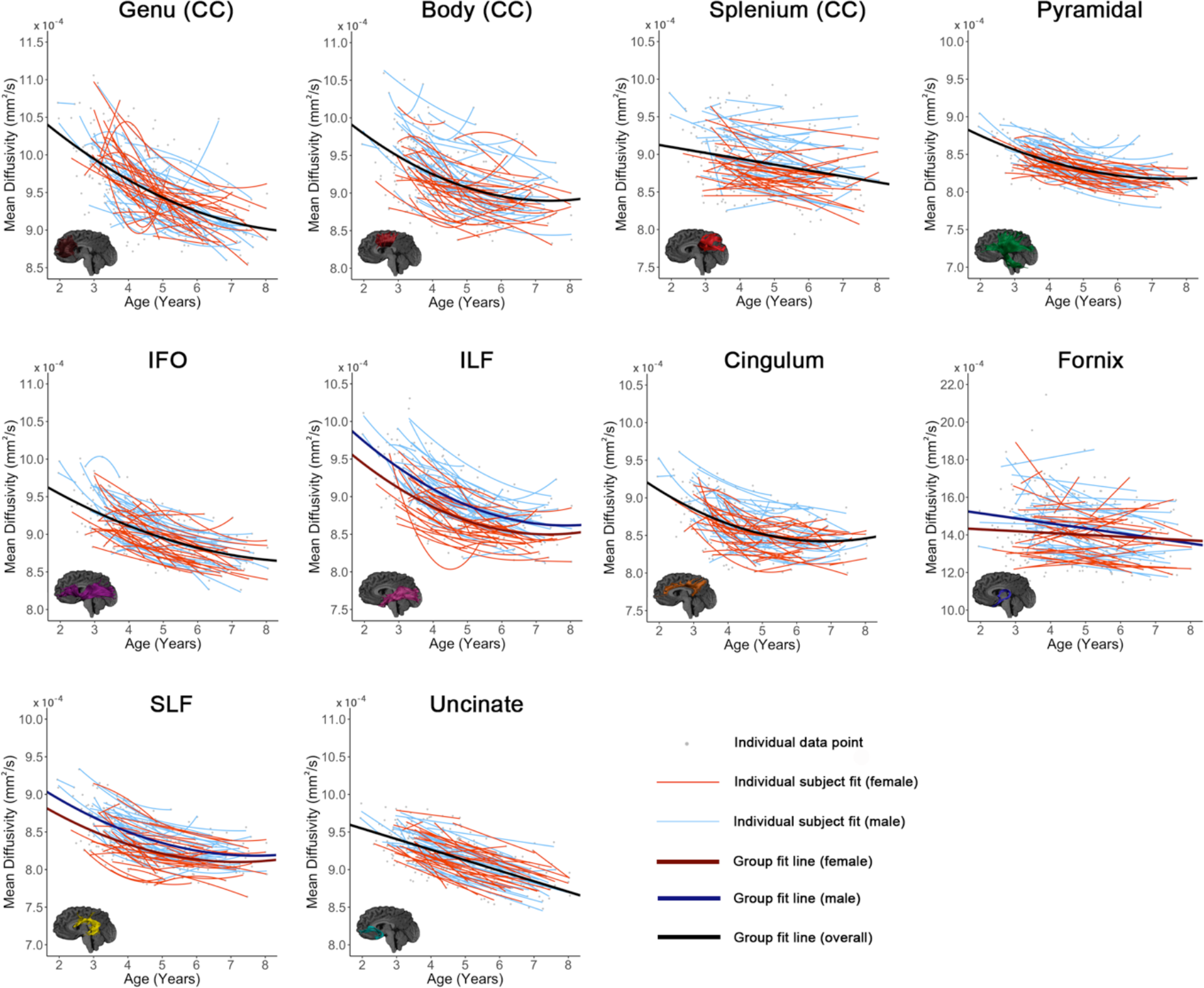
Relationships between age and MD are shown for all tracts. Best fit lines for the entire dataset are shown in black (where there was no significant age*sex interaction) or red and blue (separated for girls and boys, where there was a significant interaction). Best fit lines for each individual subject are shown in thinner red (girls) and blue (boys) lines. Individual data points are shown in grey.

Individual variability in intercepts was evident across all tracts (Figures 4-5). FA increases across the 6-year age range (2-8 years) ranged from 0.032-0.058 for the 10 tracts, equivalent to 6.1 – 14.9% change (Table 1). MD decreases ranged from 0.47*10^−4^ - 1.28*10^−4^ mm^2^/s, or 5.1 – 12.4% across individual tracts (Table 2). For FA, the splenium displayed the smallest increase, representing the slowest rate of development (i.e., the shallowest slope), while the ILF displayed the fastest increases (i.e., steepest slope). For MD, the splenium again had the lowest decrease, and the genu, followed by the ILF, had the highest decrease. For models including volumes remaining as a covariate, all model fits (e.g., linear, quadratic) remained the same as models without. For models including tract volume as a covariate, all fits remained the same as the unadjusted models, except for the genu for MD (quadratic to linear model fit), and FA for pyramidal (linear to quadratic model fit) and SLF (quadratic to linear model fit). For models including whole brain white matter volume as a covariate, all fits remained the same as those without, except for the SLF, body, and cingulum (quadratic to linear) for FA, and the genu for MD (quadratic to linear).

### 3.2 Sex differences in white matter development

The mixed models tested both a main effect of sex and age*sex interactions. A main effect of sex on FA was significant in the genu (*p* = 0.042) and pyramidal tracts (*p* = 0.005), where males had higher FA than females. Females had lower global MD than males (*p* = 0.001), and lower MD in the body (*p* = 0.005), fornix (*p* = 0.005), ILF (*p* < 0.001), and SLF (*p* < 0.001).

An interaction effect of age*sex was observed in the pyramidal tract (*p* = 0.033), with females having faster FA increases than males. A significant interaction for MD was identified in the fornix (*p* = 0.023), ILF (*p* = 0.021), and SLF (*p* = 0.021) where males had faster MD declines than females.

### 3.3 Axial and radial diffusivity changes

To further understand the processes driving FA and MD changes, we tested age related changes in axial and radial diffusivity (AD, RD). AD is more specific to axon internal structure and coherence, while RD is more specific to myelination and axonal packing (Song et al., 2005). Decreases of both AD and RD were seen with age across all tracts (Examples are shown in Figure 6; Supplementary Information Tables 1 and 2). For AD, the IFO, and uncinate had linear declines, and the cingulum, body, genu, splenium, ILF, pyramidal, and SLF had quadratic declines. Females had lower AD values than males in the body and splenium of the corpus callosum, fornix, ILF, pyramidal tract, and SLF (*p* < 0.05). Age*sex interaction effects were observed in the fornix, ILF and SLF, with males displaying faster and greater AD decreases than females (*p* <0.05). For RD, the splenium, and fornix had linear declines, and the cingulum, body, genu, IFO, ILF, pyramidal, and SLF had quadratic declines. Females had lower absolute RD intercept values than males in the body of the corpus callosum, fornix, ILF, and SLF (*p* < 0.05). Age*sex interaction effects were observed in the fornix, with males displaying faster and greater RD decreases than females (*p* <0.05)

**Figure 6.**
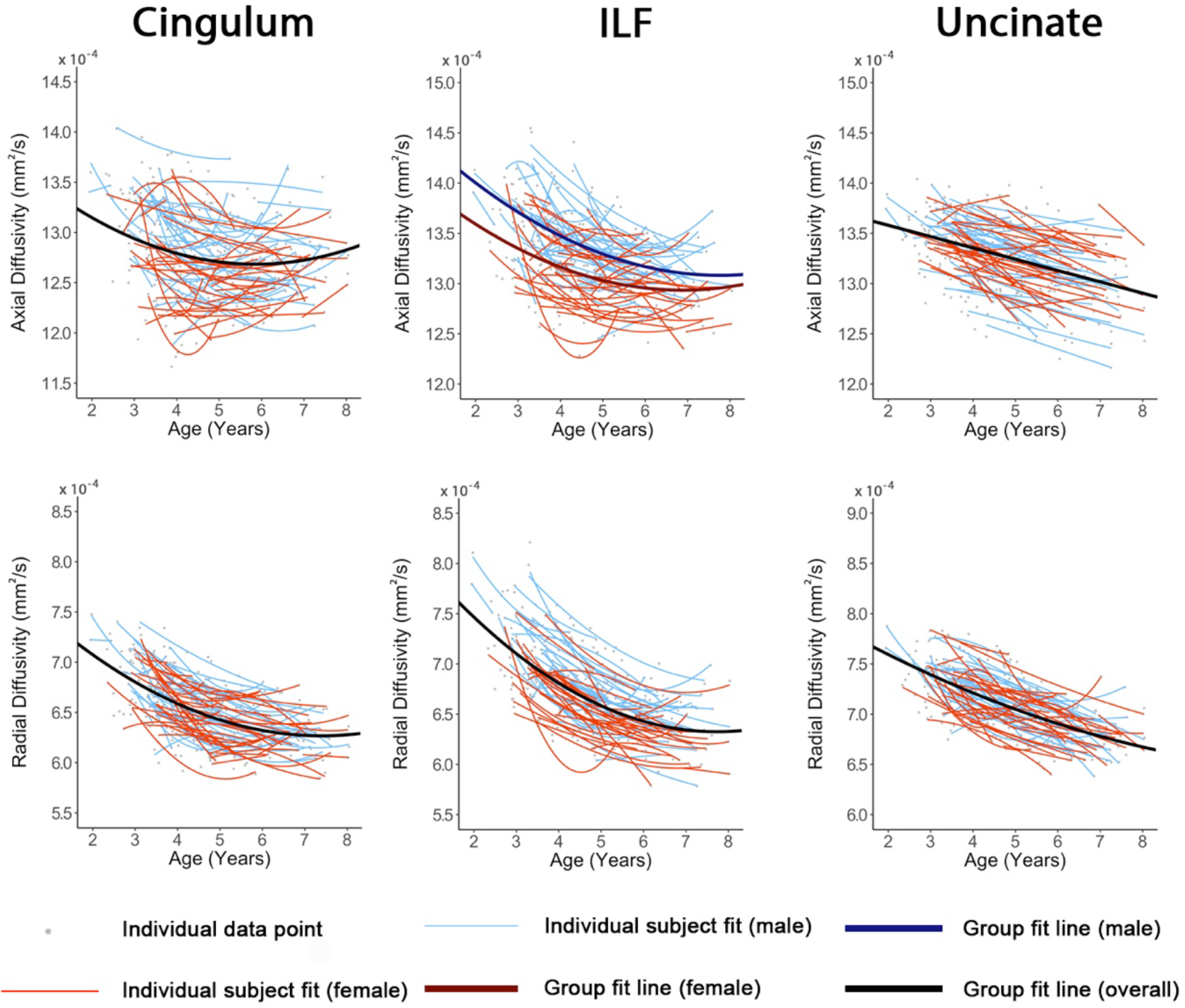
Relationships between age and axial (AD) and radial (RD) diffusivities. The cingulum, ILF, and uncinate fasciculus are presented as examples. Best fit lines for the entire dataset are shown in black (where there was no significant age*sex interaction). Best fit lines for each individual subject are shown in thinner red (girls) and blue (boys) lines. Individual data points are shown in grey.

## 4. Discussion

Our large, longitudinal study characterizes regional and temporal variation in the development of white matter connections in young children, and demonstrates earlier development in females, but faster and greater development in males during the early childhood period. Global changes of ~8.0-8.5% were observed, while changes within specific tracts ranged from 5-15%. The regional patterns suggest ongoing development of all white matter tracts, but relatively faster development within occipital and limbic tracts. Both callosal tracts and frontal-temporal association fibers have relatively slower development rates. Based on previous literature showing large changes in callosal areas in infants (Dubois et al., 2014; Paydar et al., 2014) and early maturation in adolescents (Simmonds et al., 2014), these slower changes in early childhood likely reflect the end stages of development for callosal fibers. They also are consistent with the view that development in frontal-temporal connections has not yet peaked. Considerable variation was observed in individual intercepts, perhaps suggesting intrinsic factors (e.g., genetics, prenatal environment) that influence brain structure. On the other hand, much more subtle variation is observed in development slopes, which may be used to develop a better understanding of critical understanding relationships between ongoing environmental exposures (e.g., learning) and brain development.

Different rates of change for boys and girls were observed for several tracts. In all cases, boys showed faster rates of change (steeper slopes) than females. Previous findings in older children are mixed, and many studies report no significant differences between boys and girls in rates of change (Bava et al., 2010; Bonekamp et al., 2007; Eluvathingal et al., 2007; Giorgio et al., 2010; Giorgio et al., 2008; Lebel et al., 2008b; Uda et al., 2015). A small number of studies that have tested age-sex interactions, or looked at male and female trajectories separately, consistently observed higher initial maturity in females (i.e., higher FA, lower MD) and faster rates of change in males (Asato et al., 2010; Clayden et al., 2012; Seunarine et al., 2016; Simmonds et al., 2014; Wang et al., 2012), suggesting earlier development in females. Our observations also show more substantial changes in boys. This suggests either that boys’ brains have more substantial changes across age ranges, or that the faster and earlier development in girls occurs in infancy. The literature on brain development in infants does not yet provide evidence to help interpret these findings, as most studies have not separated males and females (Dubois et al., 2014; Hermoye et al., 2006; Paydar et al., 2014). The current results point to the value of considering not just sex differences, but also how age and sex interact in studies of development.

The results of this study suggest subtle regional variations in brain development, with three different patterns emerging. Callosal and pyramidal tracts had high initial FA and slow development rates; this is most evident in the splenium of the corpus callosum. Here, the high initial FA values and low MD values suggest that these tracts are already more mature than others, and the gentler slopes imply that development is slowing down as these tracts move toward their peak values. This interpretation is consistent with infant literature, which demonstrates rapid development in callosal and pyramidal white matter (Dubois et al., 2014; Paydar et al., 2014), as well as studies in adolescence showing that these tracts are among the first to reach development plateaus (Simmonds et al., 2014). This early maturation likely reflects the fundamental role of the callosal tracts in interhemispheric communication, including motor function (Aboitiz and Montiel, 2003).

A second group of tracts has low initial FA values and shows rapid development. This is most prominently observed in the ILF, but is also apparent in in the IFO, cingulum, and fornix. It suggests that early childhood is an important period for the development of these tracts and this may be related to the rapid development of associated cognitive and behavioural skills. For example, the ILF and IFO are both heavily involved in the development of lower level visual processing and early language skills (Almairac et al., 2015; Catani et al., 2003), which show rapid development in the early childhood period. The cingulum and fornix are both limbic tracts involved in memory and spatial working memory tasks (Aggleton and Brown, 1999; Ly et al., 2016; Mishkin, 1982) and emotion, processes that continue to develop into adolescence. The ILF and IFO have been shown to have slower development and less absolute change in diffusion metrics from 5-30 years of age (Lebel et al., 2008b), which is consistent with our findings of faster development during the early childhood period. The tracts in this second group (ILF, IFO, cingulum, and fornix) also reach their peak FA/MD values earlier than other tracts (Lebel et al., 2012; Simmonds et al., 2014).

The third group of tracts, the uncinate fasciculus and SLF, had low initial FA values, like the second group, but had slower development rates. This, combined with previous literature in older children showing that these association tracts continue developing into early adulthood, reaching their peak values later than other regions, suggests they are relatively immature in the early childhood period and that that major development is yet to come (Lebel and Beaulieu, 2011; Lebel et al., 2012; Simmonds et al., 2014). The uncinate fasciculus and SLF are both frontal-temporal association tracts that underlie higher order cognitive functions, language, emotion processing, and executive function (Catani et al., 2007; Parker et al., 2005). In addition to involvement in language processing, the SLF is considered critical for general intelligence, which continues to develop over childhood and into adulthood (Jung and Haier, 2007). Both the SLF and uncinate have been implicated in psychiatric mood disorders that typically emerge during adolescence, including schizophrenia (Kubicki et al., 2007) and bipolar disorder (Heng et al., 2010), indicating that deviations from normal development during this time may represent an increased risk for mental illness (Paus et al., 2008).

Across our entire age range, both RD and AD decreased. This leads to robust decreases of MD across the age range. Furthermore, because the RD decreases are larger than the AD decreases, they drive the FA increases. This most likely reflects increasing myelination and axonal density, and decreasing membrane permeability (Beaulieu, 2002; Mukherjee et al., 2002; Qiu et al., 2008; Song et al., 2005), and is generally consistent with previous research in children (Bonekamp et al., 2007; Wang et al., 2012) and adolescents (Eluvathingal et al., 2007; Giorgio et al., 2008; Simmonds et al., 2014). Histological studies (Yakovlev and Lecours, 1967) and a large myelin water fraction study (Deoni et al., 2012) also demonstrate increases in myelination across this age range. As AD is not a specific measure and this is the first longitudinal exploration of AD in this age group, we do not have a clear understanding of the mechanisms underlying this decrease. Decreases of AD could reflect thickening of fiber diameter (Takahashi et al., 2000). Inconsistent findings for AD trajectories (decreases, no changes, and increases observed) across the adolescent period (Lebel et al., 2017) may suggest that axonal internal structure and coherence play a relatively minor role compared to other processes during development.

Only subtle differences in rates of development were observed across individuals; in general, the absolute differences in diffusion metrics persist across time points, just as one might expect children who are tall at age 2 years to remain tall throughout childhood. This may indicate a strong influence of genetics or the prenatal environment on initial diffusion parameters, which then persists across early childhood. Indeed, genetic factors account for 70-80% of white matter microstructural variation in adolescents, and 30-40% in adults (Chiang et al., 2011). Nonetheless, individual variation in slopes is apparent, and appears to be more prominent in callosal tracts and the fornix. Studies of older children show that these tracts are among the first to reach a plateau, so it may be that early childhood is an important period for the development of development for these tracts, and one in which development of these areas is more sensitive to environmental influences. In contrast, individual slopes of association tracts, where maturation continues into late adolescence or early adulthood, may be dominated in early childhood by a global development pattern. The subtle variations in individual development slopes may hold the key for understanding influences of the environment, learning, and will be important to consider alongside differences in absolute values across individuals.

Although a broad understanding of developmental trends across this period (e.g., increases in FA, decreases in MD) can be gained from extrapolation between infant and late childhood/adolescence studies, there are a number of benefits to using a longitudinal study design in this age range. Previous research highlights that the type of model fits and how well they describe the data are strongly influenced by the age range (Fjell, Walhovd et al. 2010; Lebel et al., 2017). In addition to being able to gain a more comprehensive understanding of developmental trajectories for this specific age range and the relative extent of tract development, the longitudinal design highlights subtle differences in development between individuals that would not otherwise be identified. Furthermore, results in adolescent and early adult literature point to sex differences, that already appear to be established in this older age group, however, the age at which these differences emerge remains unknown, with divergence likely occurring early in development (<2 years).

One limitation of the study was the use of non-isotropic voxels. Non-isotropic voxels can underestimate of FA in regions with crossing fibers, and fiber orientation can impact the resulting diffusion metrics (Basser et al., 2000; Oouchi et al., 2007). While the results must be interpreted with this in mind, this acquisition protocol enabled a higher in plane resolution and signal-to-noise ratio while maintaining the short acquisition time required for this population. Furthermore, the numbers of, and time between, scan points was uneven across participants. The use of linear mixed models allowed us to account for this, enabling all scans to be used in model fits.

## 5. Conclusions

In conclusion, this study shows significant white matter development across the brain during the early childhood, with faster changes in males in a number of tracts. Callosal and pyramidal tracts had high initial maturity and slow rates of change, which likely reflects progression towards a plateau. The limbic tracts (fornix, cingulum) and most association areas had low initial FA and higher rates of change, reflecting that early childhood is an important period in their development. Frontal-temporal connections showed slower rates of change and initial immaturity, which suggests considerable development is yet to come. Rates of change in all tracts likely reflect their roles in cognitive and behavioral development across this age range, and the rapid development of association and limbic areas may indicate that early childhood is an optimal time for interventions targeting the cognitive and behaviors skills associated with those areas. Variability among participants was observed in individual intercepts, but only subtle differences were observed in slopes. This may indicate that genetic mechanisms or possibly the prenatal environment determine initial diffusion values, whereas the early childhood environment may contribute to subtle variation in trajectories. Our results comprehensively map white matter development during early childhood, providing data that could be used to examine deviations from normal development that may occur in in children with various diseases, disorders, or brain injuries, which have been associated with white matter development.

## Acknowledgements

This work was supported by the Canadian Institutes of Health Research (CIHR) (funding reference numbers IHD-134090, MOP-136797, New Investigator Award to C.L) and a grant from the Alberta Children’s Hospital Foundation. J.E.R was supported by an Eyes High University of Calgary Postdoctoral Scholarship and the T. Chen Fong Postdoctoral Fellowship in Medical Imaging Science. M.N.G was supported by a University of Calgary Queen Elizabeth II Graduate Studentship award and funding provided through the Department of Paediatrcs, University of Calgary to D.D. The authors thank members of the APrON study for assistance with recruitment.

## Supplementary Material

**Table S1.**
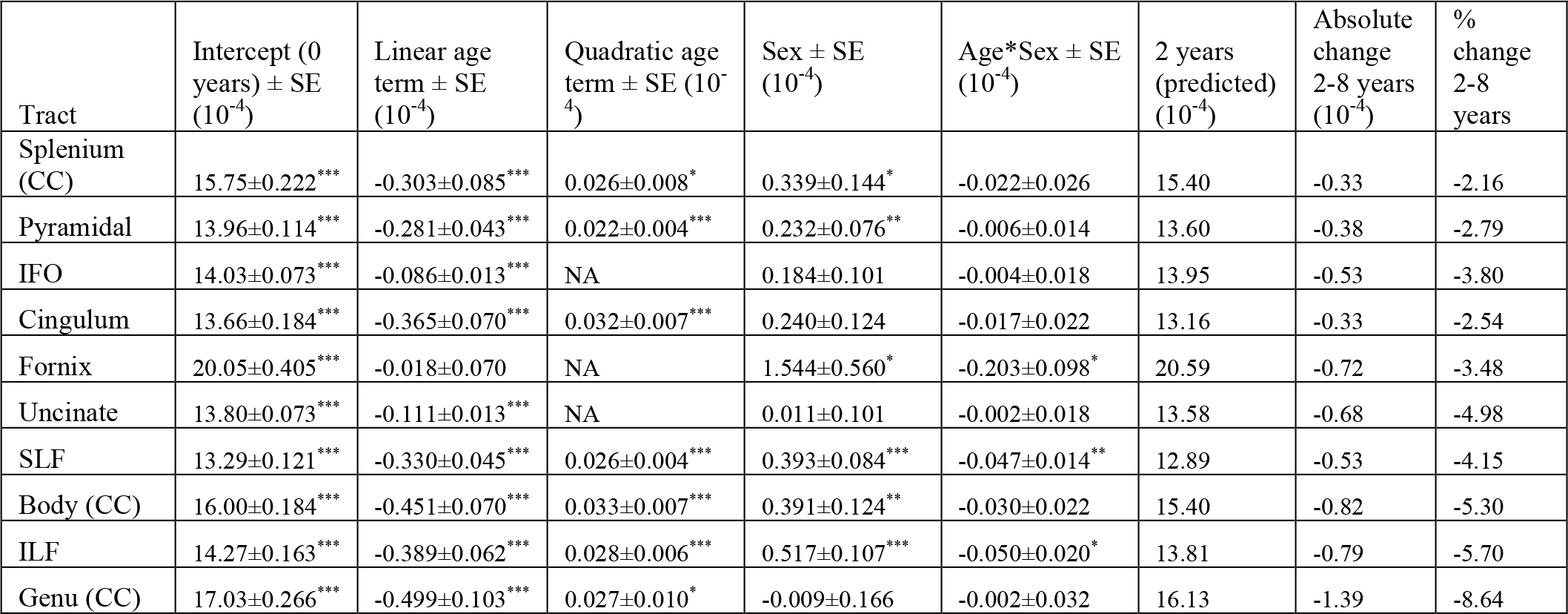
Linear regression parameters for AD (mm^2^/s) values. Tracts ordered by percent change from 2-8 years (lowest to highest). Positive sex coefficients indicate higher AD in boys. Negative age*sex interaction coefficients indicate more negative slopes in boys (i.e., faster decreases of AD). Significance levels presented as *** <0.001, ** <0.0025 (Bonferroni corrected), * <0.05.

**Table S2.** Linear regression parameters for RD (mm^2^/s) values. Tracts ordered by percent change from 2-8 years (lowest to highest). Positive sex coefficients indicate higher RD in boys. Negative age*sex interaction coefficients indicate more negative slopes in boys (i.e., faster decreases of RD). Significance levels presented as *** <0.001, ** <0.0025 (Bonferroni corrected), * <0.05.

| Tract | Intercept (0 years) ± SE (10<sup>-4</sup>) | Linear age term ± SE (10<sup>-4</sup>) | Quadratic age term ± SE (10<sup>-4</sup>) | Sex ± SE (10<sup>-4</sup>) | Age*Sex ± SE (10<sup>-4</sup>) | 2 years (predicted) (10<sup>-4</sup>) | Absolute change 2-8 years (10<sup>-4</sup>) | % change 2-8 years |
| --- | --- | --- | --- | --- | --- | --- | --- | --- |
| Splenium (CC) | 6.15±0.073*** | -0.086±0.013*** | NA | 0.148±0.101 | -0.009±0.018 | 6.05 | -0.54 | -9.01 |
| Pyramidal | 6.87±0.080*** | -0.298±0.030*** | 0.018±0.003*** | -0.035±0.054 | 0.014±0.010 | 6.34 | -0.68 | -10.79 |
| Uncinate | 8.05±0.106*** | -0.250±0.040*** | 0.010±0.004* | 0.036±0.070 | -0.004±0.013 | 7.61 | -0.93 | -12.27 |
| Fornix | 11.71±0.267*** | -0.138±0.047** | NA | 1.015±0.370* | -0.152±0.065* | 11.79 | -1.29 | -10.91 |
| SLF | 7.34±0.084*** | -0.345±0.031*** | 0.022±0.003*** | 0.194±0.061** | -0.012±0.010 | 6.82 | -0.77 | -11.35 |
| Cingulum | 7.76±0.121*** | -0.414±0.046*** | 0.028±0.004*** | 0.110±0.080 | -0.010±0.015 | 7.09 | -0.81 | -11.46 |
| Body (CC) | 7.71±0.144*** | -0.445±0.054*** | 0.030±0.005*** | 0.225±0.100* | -0.023±0.017 | 7.03 | -0.94 | -13.38 |
| Genu (CC) | 8.22±0.202*** | -0.431±0.078*** | 0.022±0.008* | -0.188±0.128 | 0.016±0.024 | 7.37 | -1.21 | -16.46 |
| ILF | 8.30±0.115*** | -0.537±0.043*** | 0.035±0.004*** | 0.294±0.079*** | -0.023±0.014 | 7.49 | -1.16 | -15.49 |
| IFO | 7.93±0.098*** | -0.372±0.037*** | 0.020±0.004*** | 0.093±0.067 | -0.005±0.012 | 7.31 | -1.06 | -14.48 |

**Table S3.**
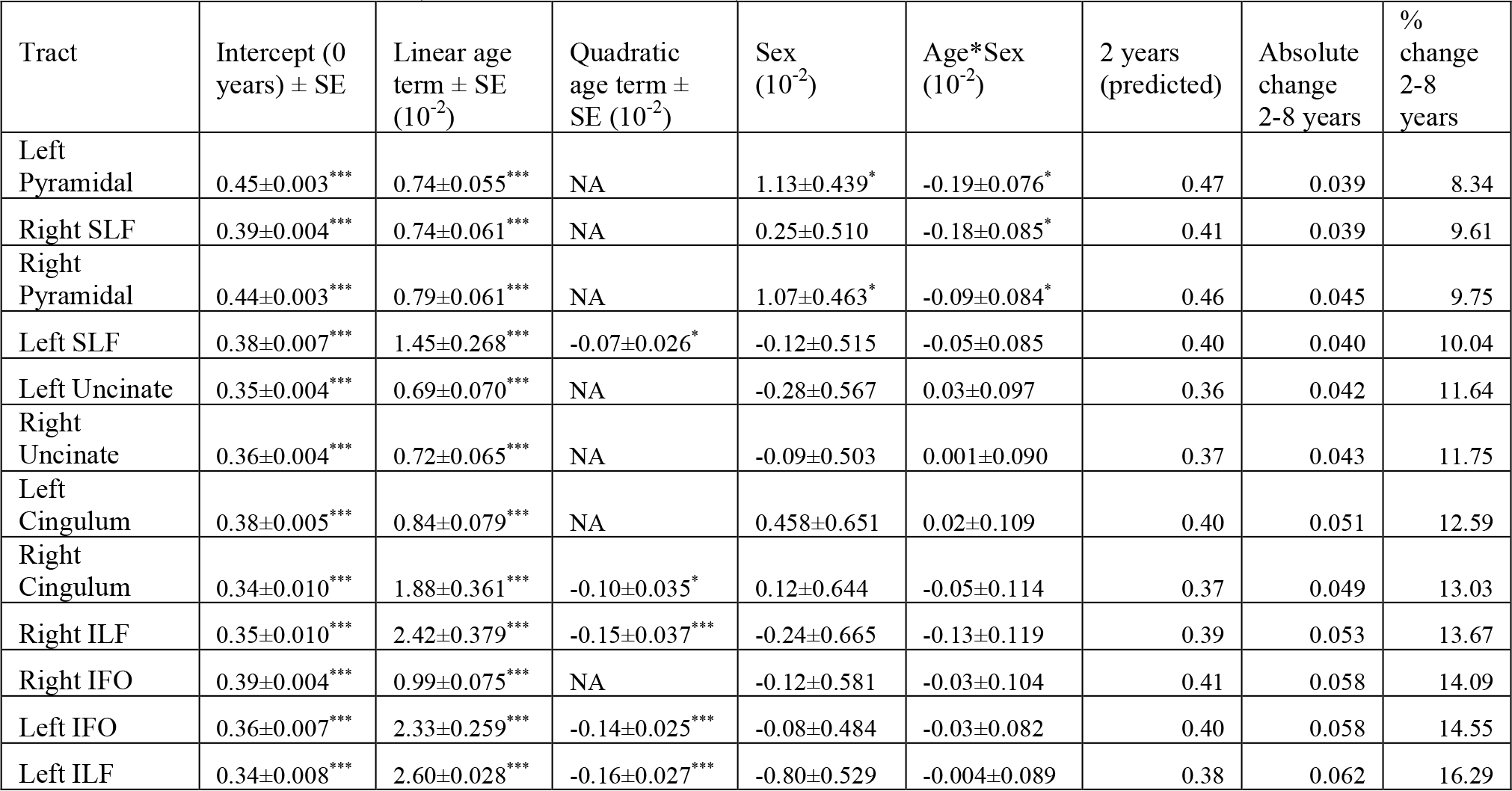
Linear regression parameters for FA for left and right hemisphere tracts. Tracts are ordered by percent change from 2-8 years (lowest to highest). Sex term represents boys: Positive sex coefficients indicate higher FA in boys; Positive age*sex interaction coefficients indicate more positive slopes in boys (i.e., faster increases of FA). Significance levels presented as *** <0.001, ** <0.0025 (Bonferroni corrected), * <0.05.

**Table S4.** Linear regression parameters for MD (mm^2^/s) for left and right hemisphere tracts. Tracts ordered by percent change from 2-8 years (lowest to highest). Sex term represents boys: Positive sex coefficients indicate higher MD in boys; Negative age*sex interaction coefficients indicate more negative slopes in boys (i.e., faster decreases of MD). Significance levels presented as *** <0.001, ** <0.0025 (Bonferroni corrected), * <0.05.

| Tract | Intercept (0 years) ± SE (10<sup>-4</sup>) | Linear age term ± SE (10<sup>-4</sup>) | Quadratic age term ± SE (10<sup>-4</sup>) | Sex ± SE (10<sup>-4</sup>) | Age*Sex ± SE (10<sup>-4</sup>) | 2 years (predicted) (10<sup>-4</sup>) | Absolute change 2-8 years (10<sup>-4</sup>) | % change 2-8 years |
| --- | --- | --- | --- | --- | --- | --- | --- | --- |
| Left Pyramidal | 9.25±0.091*** | -0.309±0.035*** | 0.021±0.003*** | 0.079±0.060 | 0.005±0.011 | 8.76 | -0.57 | -6.45 |
| Right Pyramidal | 9.22±0.098*** | -0.276±0.038*** | 0.017±0.004*** | 0.039±0.064 | 0.007±0.012 | 8.76 | -0.60 | -6.85 |
| Right SLF | 9.27±0.109*** | -0.333±0.041*** | 0.024±0.004*** | 0.245±0.074** | -0.020±0.013 | 8.80 | -0.62 | -7.00 |
| Left Cingulum | 9.76±0.128*** | -0.387±0.049*** | 0.028±0.005*** | 0.129±0.083 | -0.013±0.015 | 9.15 | -0.65 | -7.14 |
| Right Cingulum | 9.70±0.143*** | -0.412±0.055*** | 0.031±0.005*** | 0.185±0.092* | -0.013±0.017 | 9.08 | -0.66 | -7.24 |
| Right Uncinate | 9.69±0.056*** | -0.130±0.011*** | NA | -0.002±0.077 | 0.002±0.015 | 9.43 | -0.77 | -8.22 |
| Left SLF | 9.38±0.099*** | -0.348±0.037*** | 0.023±0.004*** | 0.276±0.068*** | -0.027±0.012* | 8.89 | -0.77 | -8.67 |
| Right IFO | 9.62±0.070*** | -0.141±0.013*** | NA | 0.109±0.097 | 0.001±0.018 | 9.39 | -0.84 | -8.96 |
| Left Uncinate | 9.92±0.061*** | -0.143±0.011*** | NA | 0.085±0.084 | -0.015±0.016 | 9.66 | -0.90 | -9.34 |
| Left IFO | 10.21±0.120*** | -0.366±0.046*** | 0.022±0.004*** | 0.147±0.079 | -0.013±0.014 | 9.63 | -0.94 | -9.72 |
| Right ILF | 10.07±0.168*** | -0.439±0.065*** | 0.030±0.006*** | -0.379±0.108*** | -0.039±0.020 | 9.46 | -0.92 | -9.77 |
| Left ILF | 10.52±0.138*** | -0.539±0.053*** | 0.036±0.005*** | -0.346±0.092*** | -0.021±0.017 | 9.74 | -1.16 | -11.88 |

**Figure S1.**
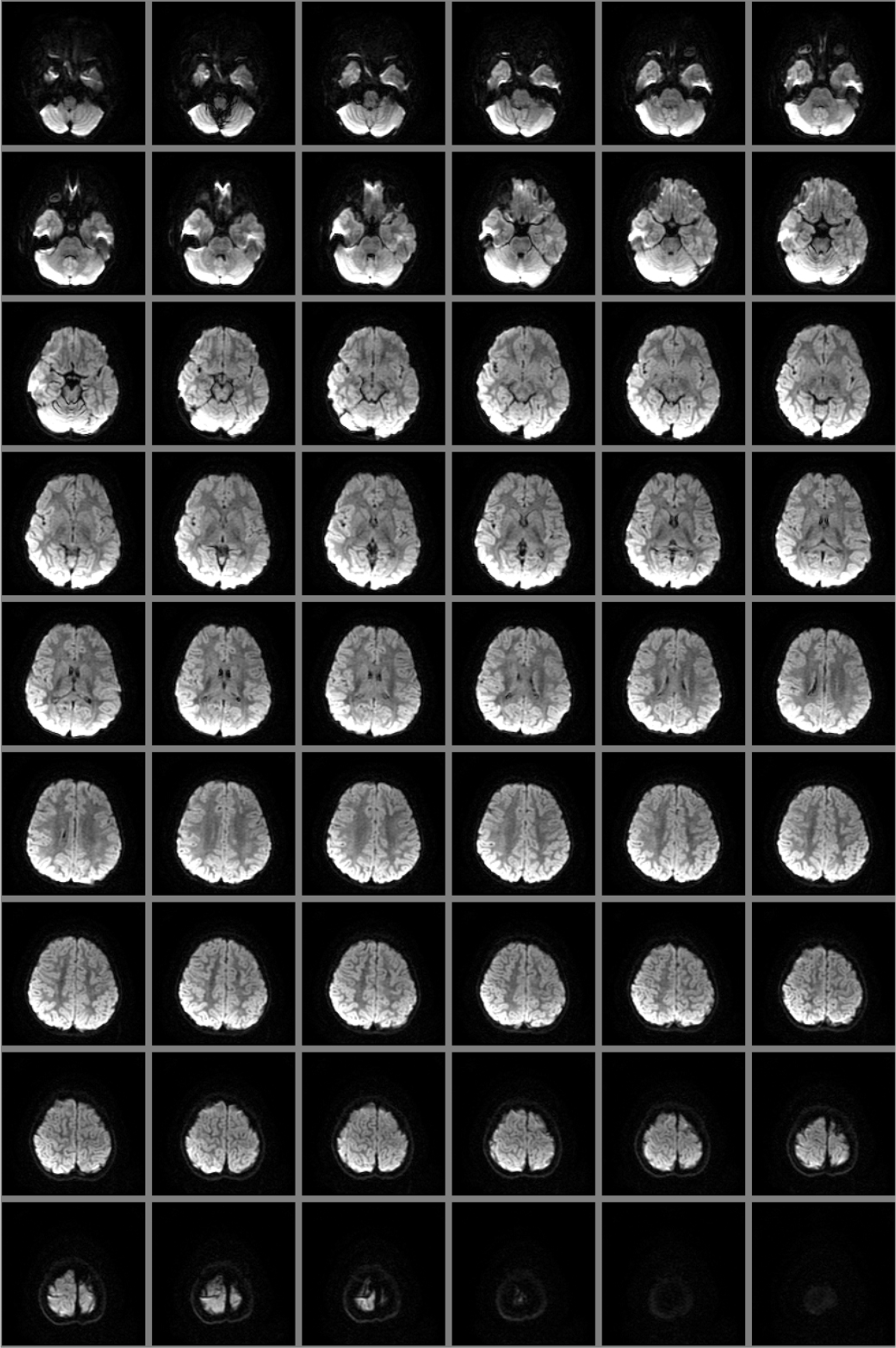
Representative diffusion weighted image (axial slices) from a 4.64-year-old female.

**Figure S2.**
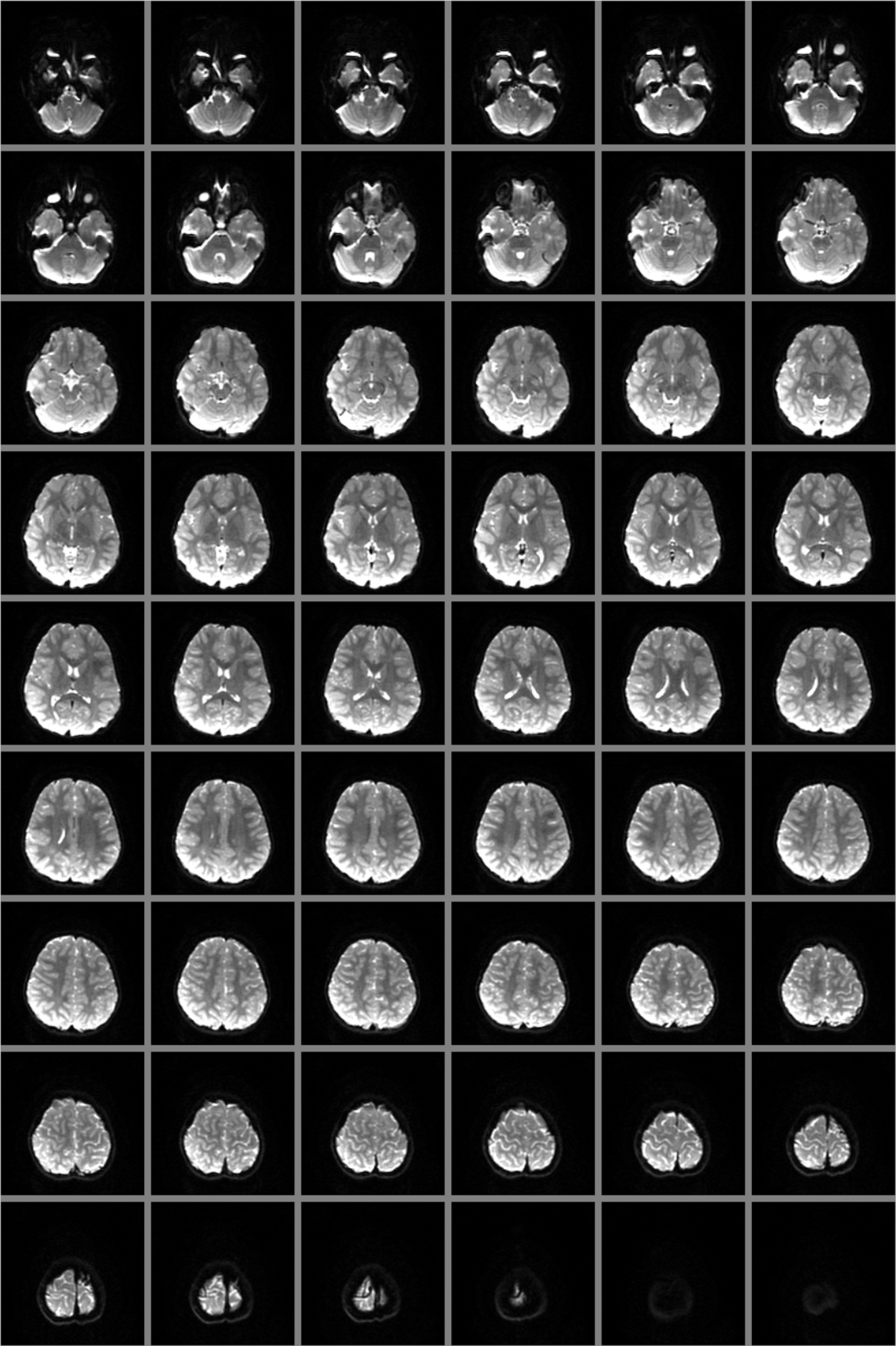
Representative b0 image (axial slices) from a 4.64-year-old female.

**Figure S3.**
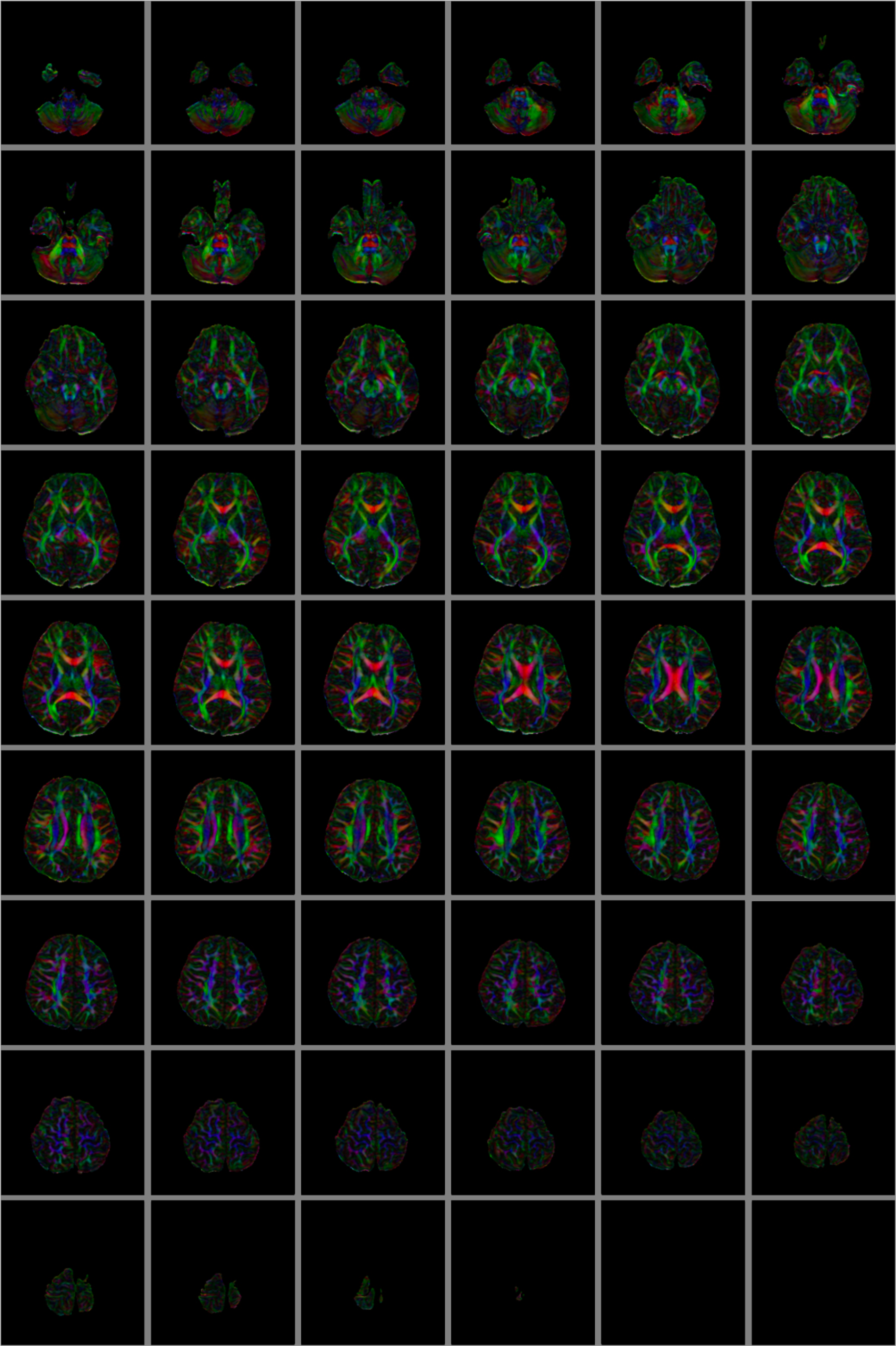
Representative color map (axial slices) from processed data from a 4.64-year-old female.

**Figure S4.**
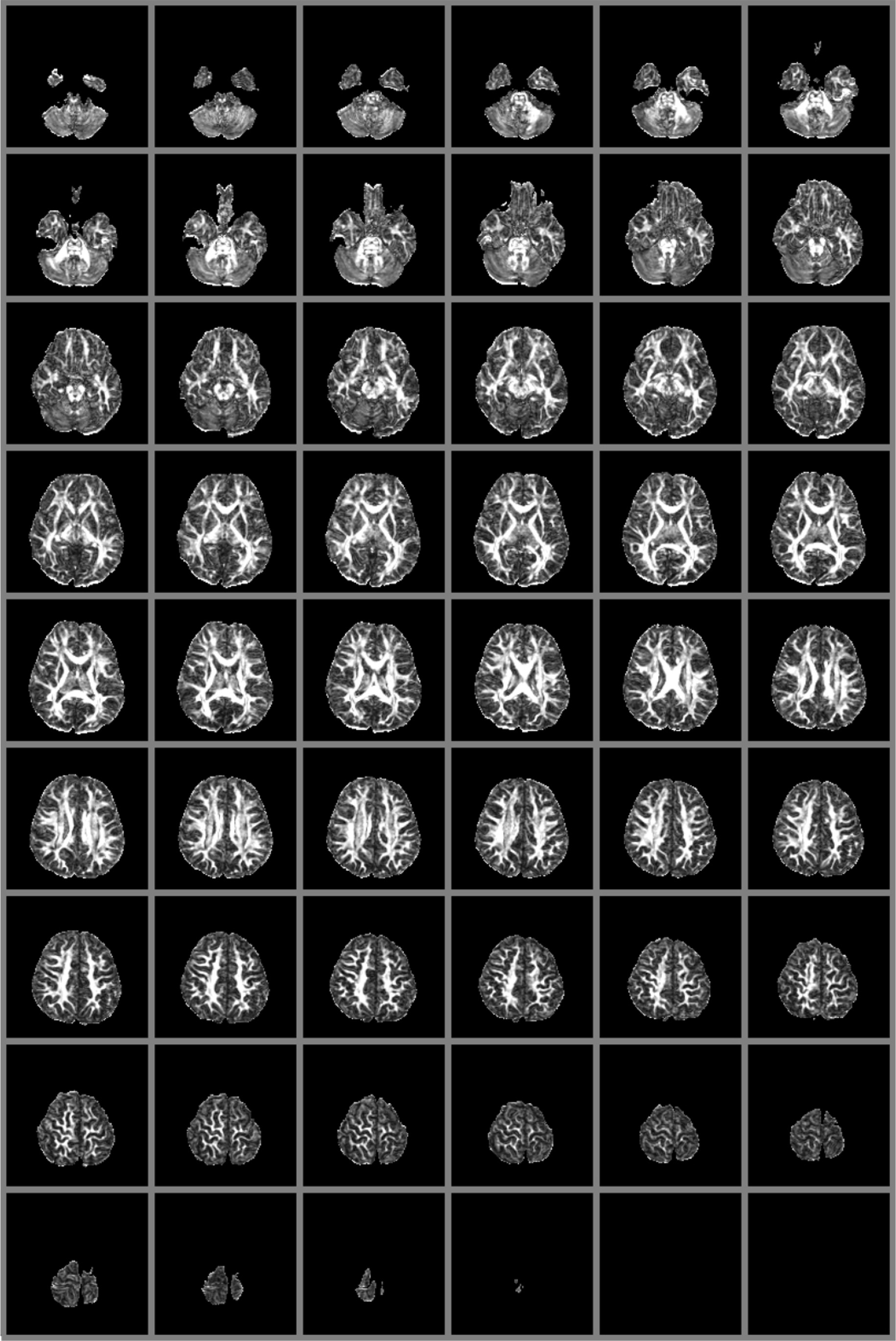
Representative FA map (axial slices) from processed data from a 4.64-year-old female.

**Figure S5.**
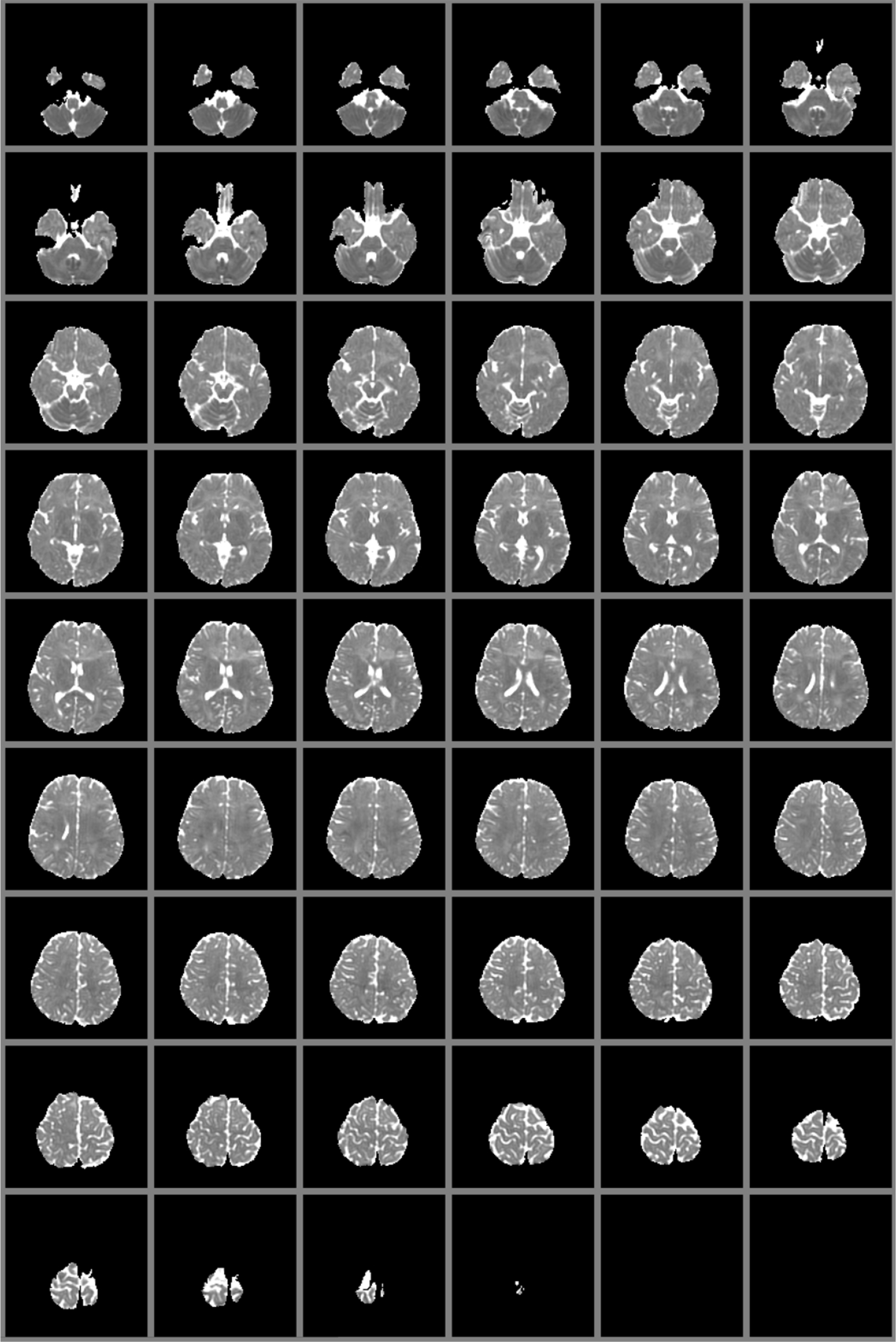
Representative MD map (axial slices) from processed data from a 4.64-year-old female.

**Figure S6.**
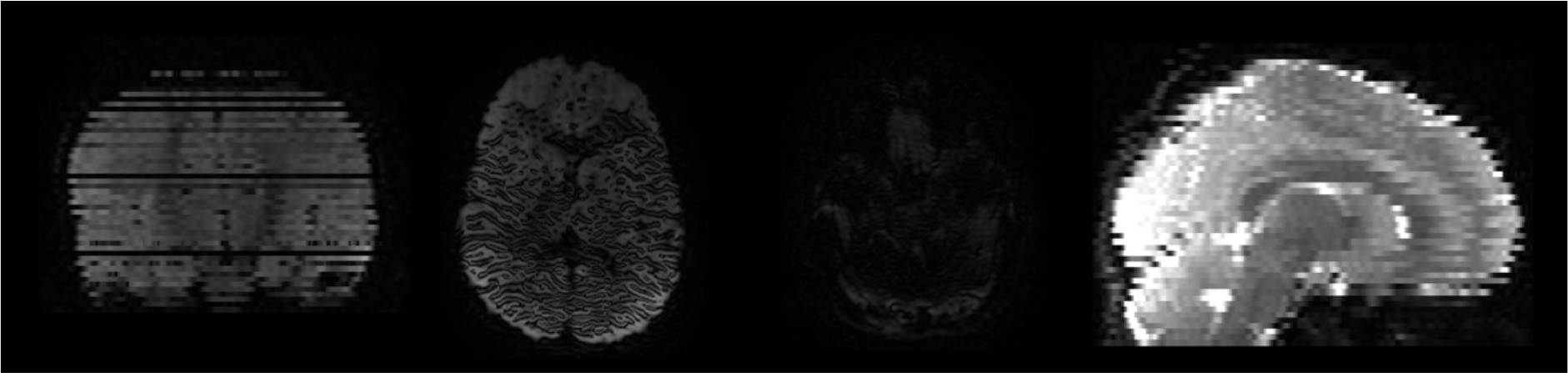
Examples of the most commonly identified artifacts in the dataset.

## Notes

**Conflicting of Interest** The authors report no conflicting interests.

